# Testing Causal Bidirectional Influences between Physical Activity and Depression using Mendelian Randomization

**DOI:** 10.1101/364232

**Authors:** Karmel W. Choi, Chia-Yen Chen, Murray B. Stein, Yann C. Klimentidis, Min-Jung Wang, Major Depressive Disorder Working Group of the Psychiatric Genomics Consortium, Karestan C. Koenen, Jordan W. Smoller

## Abstract

**Background:** Burgeoning evidence from randomized controlled trials and prospective cohort studies suggests that physical activity protects against depression, pointing to a potential modifiable target for prevention. However, the direction of this inverse association is not clear: physical activity may reduce risk for depression, and/or depression may result in decreased physical activity. Here, we used bidirectional two-sample Mendelian randomization (MR) to test causal influences between physical activity and depression.

**Methods:** For genetic instruments, we selected independent top SNPs associated with major depressive disorder (MDD, N = 143,265) and two physical activity phenotypes—self-reported (N = 377,234) and objective accelerometer-based (N = 91,084)—from the largest available, non-overlapping genome-wide association results. We used two sets of genetic instruments: (1) only SNPs previously reported as genome-wide significant, and (2) top SNPs meeting a more relaxed threshold (p < 1×10^-7^). For each direction of influence, we combined the MR effect estimates from each instrument SNP using inverse variance weighted (IVW) meta-analysis, along with other standard MR methods such as weighted median, MR-Egger, and MR-PRESSO.

**Results:** We found evidence for protective influences of accelerometer-based activity on MDD (IVW odds ratio (OR) = 0.74 for MDD per 1 SD unit increase in average acceleration, 95% confidence interval (CI) = 0.59-0.92, p =.006) when using SNPs meeting the relaxed threshold (i.e., 10 versus only 2 genome-wide significant SNPs, which provided insufficient data for sensitivity analyses). In contrast, we found no evidence for negative influences of MDD on accelerometer-based activity (IVW b = 0.04 change in average acceleration for MDD versus control status, 95% CI = −0.43-0.51, p =.87). Furthermore, we did not see evidence for causal influences between self-reported activity and MDD, in either direction and regardless of instrument SNP criteria.

**Discussion:** We apply MR for the first time to examine causal influences between physical activity and MDD. We discover that objectively measured—but not self-reported—physical activity is inversely associated with MDD. Of note, prior work has shown that accelerometer-based physical activity is more heritable than self-reported activity, in addition to being more representative of actual movement. Our findings validate physical activity as a protective factor for MDD and point to the importance of objective measurement of physical activity in epidemiological studies in relation to mental health. Overall, this study supports the hypothesis that enhancing physical activity is an effective prevention strategy for depression.

## Introduction

Depression is a common psychiatric condition that represents a leading cause of disability and lost productivity worldwide (1). Despite this, efforts to prevent depression have been challenging, with few established protective factors, particularly modifiable targets for prevention. One promising modifiable target is physical activity, defined broadly as musculoskeletal movement resulting in energy expenditure (2). The relationship between physical activity and depression has received much attention in recent years, with evidence suggesting protective effects of physical activity on depression. For example, meta-analytic studies of randomized controlled trials (RCTs) have shown that physical activity is associated with moderate but notable reductions of depressive symptoms in at-risk populations (3), and prospective studies have demonstrated associations between higher physical activity and decreased risk for later depression (4,5).

While such findings point to a protective role of physical activity for depression, several questions remain. First, to what extent is the relationship between physical activity and depression bidirectional? Some studies have suggested that depression is associated with reduced physical activity (6,7), but few studies have simultaneously tested both directional effects. Second, does measurement of physical activity matter? Most studies to date have relied on self-reported measures of activity, which may be subject to mood effects, memory inaccuracy, and social desirability bias (8). In addition, RCTs often assume that treatment assignment is equivalent to increased physical activity, without necessarily measuring actual changes in physical activity. Third, is the observed protective effect of physical activity on depression free of potential confounding? Although RCTs minimize bias and confounders by design, they are intensive to conduct and have been of relatively limited size. Further approaches are needed to address these limitations.

Mendelian randomization (MR) is a method that leverages the random assortment of genetic variants (i.e., single nucleotide polymorphisms, or SNPs) prior to birth to establish instruments for causal inference free of usual sources of environmental or genetic confounding. The premise of MR (Figure 1) is that SNPs strongly associated with a non-genetic exposure of interest (e.g., physical activity) can be used as proxies for that exposure to test putatively causal effects on the outcome (e.g., depression). MR assumes that exposure-associated SNPs are not directly related with the outcome itself, nor with other variables that cause the outcome through a different pathway (i.e., horizontal pleiotropy). In a two-sample MR design, SNPs can be extracted from summary statistics of large-scale genome-wide association studies (GWAS), which have recently become available for physical activity (9) and major depressive disorder (10). Here, we apply bidirectional MR to test causal effects of physical activity on depression, and vice versa. Furthermore, we examine genetic instruments based on self-reported versus objectively measured activity.

**Figure 1.**
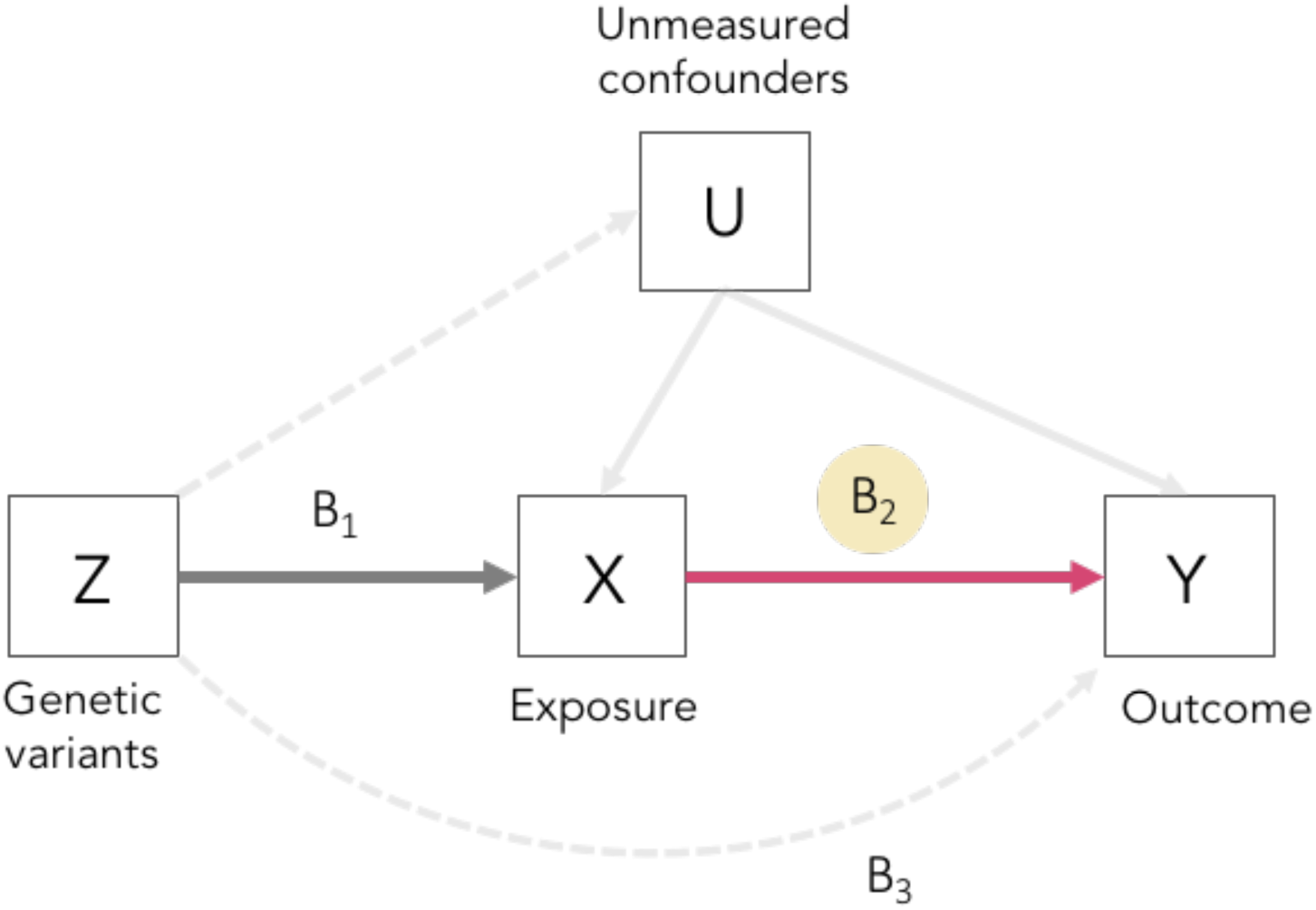
Mendelian randomization (MR) model. B1 and B3 represent estimated direct effects of exposure-related genetic variants on the exposure (e.g., physical activity) and the outcome (e.g., depression); B2 is the causal effect of interest to be estimated. Solid paths are theorized to exist; dashed paths are theorized to be non-significant according to **MR** assumptions.

## Methods

### Data sources and instruments

#### Physical activity

We drew on summary statistics from a recent GWAS of physical activity conducted in the UK Biobank (9). This GWAS examined two continuous activity phenotypes: (1) self-reported moderate-to-vigorous physical activity (MVPA; in inverse-normalized metabolic-equivalent minutes per week); and (2) objective accelerometer-based activity, specifically overall acceleration average (in milligravities across at least 72 hours of accelerometer wear). The GWAS for self-reported activity (N = 377,234) identified nine independent genome-wide significant (GWS) SNPs, though SNP-based heritability was modest at around 5% (9). The GWAS for accelerometer-based activity (N = 91,084) identified only two independent GWS SNPs, though SNP-based heritability was estimated much higher at 14%. These heritability estimates indicate that SNPs beyond those currently identified as GWS may contribute to variation in physical activity. Given this, we used two sets of genetic instruments: (1) only SNPs previously reported as genome-wide significant (GWS; p < 5×10^-9^), and (2) top SNPs meeting a more relaxed threshold (p < 1×10^-7^). This method of relaxing the statistical threshold for genetic instruments has previously been used in psychiatric research when few GWS SNPs are available (11,12). When the more relaxed threshold was used, we clumped SNPs at r^2^ >.001 for independence based on European ancestry reference data from the 1000 Genomes Project. Where SNPs for the exposure phenotype were not available in the summary statistics of the outcome phenotype, we identified overlapping proxy SNPs in high linkage disequilibrium (r^2^>.80) using the LDproxy search on the online platform LDlink (https://analysistools.nci.nih.gov/LDlink). For resulting lists of instrument SNPs for each phenotype, see Supplementary Tables S1A-D.

#### Depression

We drew on summary statistics from the largest and most recent GWAS for major depressive disorder (MDD) (10). Overall, this case-control GWAS identified 44 independent GWS (p < 5×10^-8^) SNPs for MDD. For the MR analysis, we used meta-analytic results for MDD leaving out UK Biobank samples since the physical activity GWAS was also conducted in the UK Biobank, and also without 23andMe samples due to general access constraints. This resulted in a meta-analytic subsample of N = 143,265. As instruments, we used independent clumped SNPs meeting a relaxed threshold (p < 1×10^-6^) to account for the reduced meta-analytic subsample, with similar procedures for identifying proxy SNPs as needed. For resulting list of instrument SNPs, see Supplementary Table S1E.

### Statistical analyses

MR analyses were conducted in R using the *TwoSampleMR* package. This package harmonizes exposure and outcome datasets containing information on SNPs, alleles, effect sizes (odds ratios converted to betas by taking the log transformation), standard errors, p-values, and effect allele frequencies for selected exposure instruments. Here, we allowed the forward strand of ambiguous SNPs to be inferred where possible based on allele frequency information; however, strand ambiguous SNPs with intermediate effect allele frequencies were considered unresolvable. We also conducted sensitivity analyses where all strand ambiguous SNPs were excluded from MR analysis, which did not change the pattern of findings; thus, full results are reported.

For each direction of potential influence, we combined MR effect estimates using inverse variance weighted (IVW) meta-analysis, along with other standard MR methods such as weighted median (WM) and MR Egger, which have been shown to be more robust to invalid instruments though with reduced statistical power (13,14). We further applied MR-PRESSO (15) to detect and remove any likely outliers reflecting horizontal pleiotropy for all reported results. Finally, for any significant results we conducted further analyses to assess robustness, via leave-one-SNP-out analyses and intercept tests of horizontal pleiotropy based on deviation from the null. We further manually looked up each SNP and their proxies (r^2^ =.80) in the PhenoScanner GWAS database (http://phenoscanner.medschl.cam.ac.uk) to assess for any notable previous associations (p < 1×10^-5^) with potential confounding traits, and assessed effects of removing these SNPs.

## Results

### Accelerometer-based physical activity and depression

We found evidence for causal influences of accelerometer-based activity on MDD (IVW odds ratio (OR) = 0.74 for MDD per 1 SD unit increase in average acceleration, 95% confidence interval (CI) = 0.59-0.92, p =.006; WM and MR Egger yielded similar results—see Table 1) with 10 SNPs meeting the more liberal statistical threshold (Figure 2). The MR effect was not significant with only two GWS SNPs (IVW OR = 1.01, 95% CI = 0.96-1.07, p =.60), which also provided insufficient data for alternative MR methods and sensitivity analyses. For the 10 SNPs, MR-PRESSO did not detect any potential outliers. Furthermore, analyses leaving out each SNP revealed that no single SNP drove these results but rather reflected an overall combined pattern of opposite effects on physical activity versus MDD (Supplementary Figure S2). The MR Egger intercept test suggested no horizontal pleiotropy (intercept = 0.008, standard error = 0.02, p =.60). In the PhenoScanner database, we identified two of the 10 SNPs previously associated with potentially depression-relevant traits (i.e., rs59499656 with BMI and rs9293503 with educational attainment). However, removing both SNPs did not change the pattern of results.

**Table 1.** Mendelian randomization results of accelerometer-based activity → depression.

| Method | Odds ratio | 95% CI | p-value | SNPs |
| --- | --- | --- | --- | --- |
| Inverse-variance weighted<sup>†</sup> | 0.74 | 0.59, 0.92 | 0.006** | 10 |
| Weighted median<sup>†</sup> | 0.71 | 0.53, 0.95 | 0.02* | 10 |
| MR Egger<sup>†</sup> | 0.57 | 0.22, 1.48 | 0.28 | 10 |

\*\* p < 0.01, \* p < 0.05.
<sup>†</sup>No MR-PRESSO outliers detected. Top SNPs p < 1x10<sup>-7</sup>.
Odds ratio = risk for MDD per 1 standard deviation increase in average acceleration.

**Figure 2.**
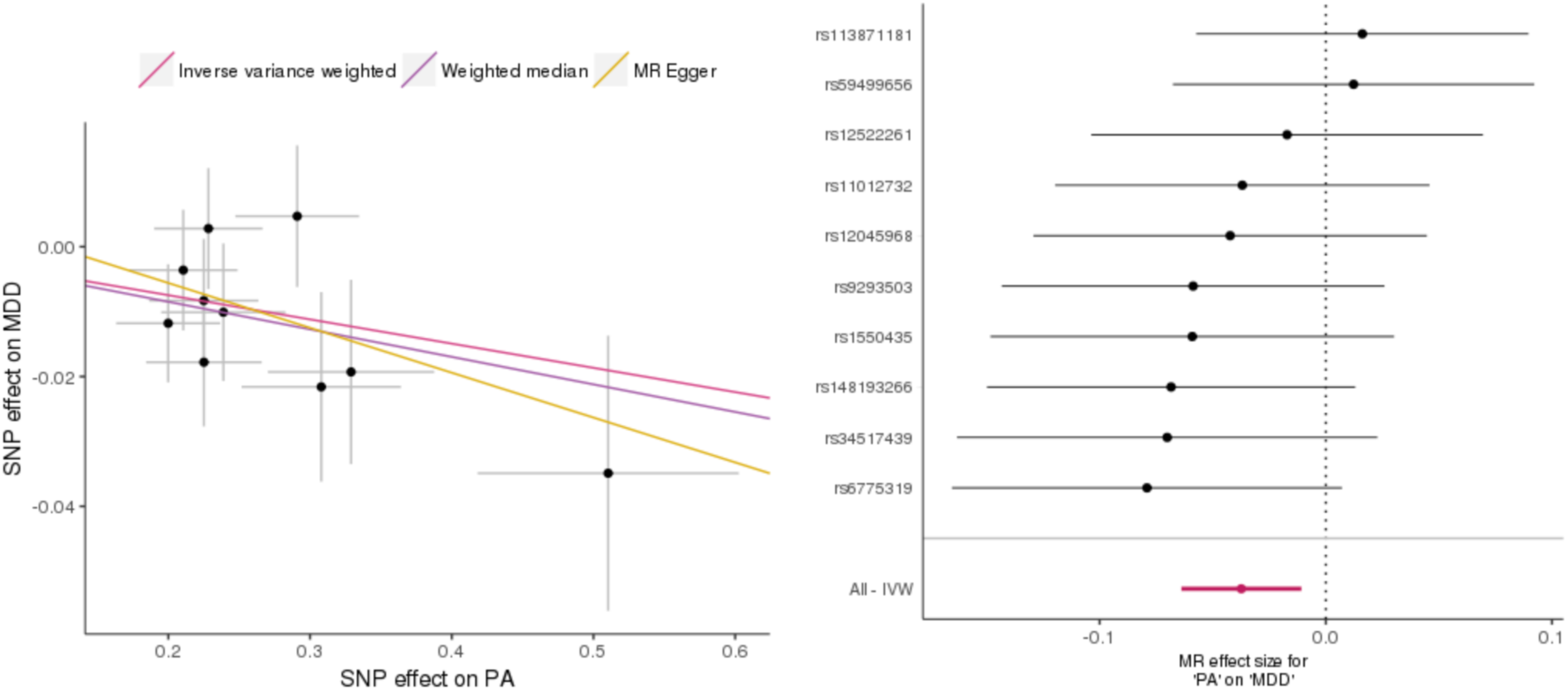
Mendelian randomization plots for accelerometer-based activity on depression. (a) Scatterplot of SNP effects on physical activity (PA) versus their effects on depression (MDD), with slope of each line corresponding to estimated MR effect per method. (b) Forest plot of individual and combined SNP effects. Top SNPs p < 1×10^-7^.

In the other direction, we found no evidence for causal influences of MDD on accelerometer-based PA, across all MR methods (Table 2). MR-PRESSO detected one outlier, and effects remained null after removal of this outlier (IVW b = 0.09 decrease in average acceleration per MDD caseness, 95% CI = −0.50-0.32, p =.66; WM and MR Egger yielded similar results—see Table 2) (Figure 3).

**Table 2.** Mendelian randomization results of depression → accelerometer-based activity.

| Method | Beta | 95% CI | p-value | SNPs |
| --- | --- | --- | --- | --- |
| Inverse-variance weighted<sup>†</sup> | -0.09 | -0.50, 0.32 | 0.66 | 15 |
| Weighted median<sup>†</sup> | -0.06 | -0.62, 0.51 | 0.83 | 15 |
| MR Egger<sup>†</sup> | -0.01 | -2.03, 2.01 | 0.99 | 15 |
| Inverse-variance weighted | 0.04 | -0.43, 0.51 | 0.87 | 16 |
| Weighted median | -0.01 | -0.56, 0.53 | 0.97 | 16 |
| MR Egger | 1.14 | -0.88, 3.16 | 0.29 | 16 |

<sup>†</sup>With MR-PRESSO outlier (rs78676209) removed. Top SNPs p < 1x10<sup>-6</sup>.
Beta = change in average acceleration with MDD caseness.

**Figure 3.**
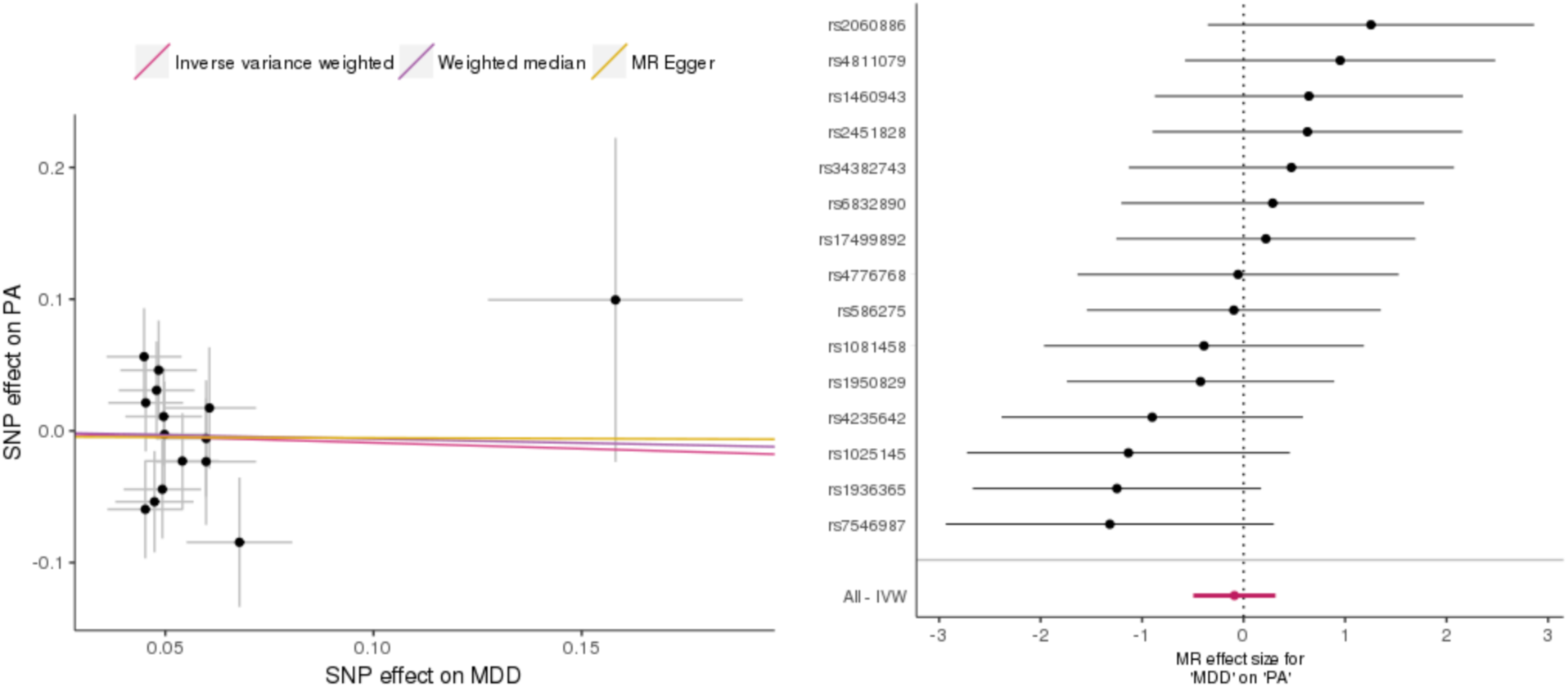
Mendelian randomization plots for depression on accelerometer-based activity. (a) Scatterplot of SNP effects on depression (MDD) versus their effects on physical activity (PA), with slope of each line corresponding to estimated MR effect per method. (b) Forest plot of individual and combined SNP effects.

### Self-reported physical activity and depression

In contrast, we found no evidence for causal influences of self-reported activity on MDD, regardless of instrument SNP threshold (outlier-adjusted IVW OR = 1.28 per 1 SD increase in average acceleration, 95% CI = 0.87-1.90, p = 0.21, for 24 top SNPs; IVW OR = 1.45 per 1 SD unit increase in average acceleration, 95% CI = 0.52-4.04, p = 0.48, for six GWS SNPs; see Supplementary Table S5 and Supplementary Figure S5), and for causal influences of MDD on self-reported activity (outlier-adjusted IVW beta = 0.02 per MDD caseness, 95% CI = −0.008-0.05, p = 0.15, for 14 top SNPs; see Supplementary Table S6 and Supplementary Figure S6).

## Discussion

Depression is a highly common and debilitating condition, with a high societal burden of morbidity and mortality (16). As such, the identification of effective strategies for preventing depression has substantial implications for improving population health. Recent evidence has suggested that physical activity may protect against risk for depression (3–5). However, if the relationship between physical activity and depression is not causal, recommendations to promote physical activity, while beneficial for other health outcomes, would yield limited results for depression. To strengthen causal inference about the effects of physical activity on depression, we applied a powerful, genetically-informed strategy: Mendelian randomization. Using genetic instruments selected from large-scale existing GWAS data, we find evidence supporting causal protective effects of physical activity on depression.

Our results extend the current literature in a number of ways. First, we examine causal effects of both self-reported and objectively measured (i.e., accelerometer-based) physical activity. We find that objectively measured—but not self-reported—physical activity is associated with reduced risk of depression. Of note, prior work has shown that objectively measured physical activity is more heritable than self-reported activity (9), in addition to being more representative of actual activity (8). Only one prior study to our knowledge has incorporated genetic information, using a twin-based design, to test causal effects of exercise on depression (17). Contrary to our present study, it did not yield evidence for any causal protective effects, perhaps owing to reliance on self-report measures and definition of physical activity based on leisure exercise, i.e., physical activity performed to improve or maintain fitness.

Second, to date it has remained unclear whether the inverse association between physical activity and depression is due to the protective effect of physical activity on depression and/or the decreasing effect of depression on physical activity. Using bidirectional MR, we test both explanations and find evidence for only one direction of causal effect, where physical activity influences depression, while depression does not appear to causally influence physical activity. Given prior evidence of associations between depression and reduced physical activity (7), our results suggest that other factors may better explain this relationship rather than depression directly influencing physical activity. For example, chronic pain can both interfere with physical activity and lead to depression.

This study has several limitations. First, while we drew on the largest available GWAS, some identified very few genome-wide significant SNPs, which may result in relatively weak genetic instruments. To address this, we applied statistical criteria to include a larger number of associated SNPs as instruments. This approach has been used in other MR research in psychiatry where available genome-wide significant SNPs are limited (11,12). Second, despite selecting strongly associated SNPs, it is known that common SNPs do not yet explain much total variance in complex traits (18)—so they cannot be considered exact proxies of the exposure itself. In addition, since we do not yet know the biological action of these SNPs, it is impossible to fully rule out pleiotropic mechanisms, though we conducted multiple sensitivity analyses for horizontal pleiotropy. We view our application of MR as a test of whether genetic instruments can provide independent support for the protective effect of physical activity on depression that has been observed in RCTs and prospective studies. Our novel contribution to this literature is the use of genetic variants as instruments for causal inference, which obviate some of the typical challenges in observational research and do recapitulate existing evidence. Third, we used summary GWAS data for MDD status and not for depressive symptoms in individuals with or without MDD. While meta-analyses have shown that physical activity can improve symptoms in depressed individuals (19–21), we were not able to address this issue. Furthermore, SNPs associated with physical activity were identified in the UK Biobank, which comprises individuals between 40 and 70 years old, whereas samples in the MDD GWAS included a wider range of age groups. It is possible that physical activity in younger individuals is influenced by other variants that share different associations with MDD, though such GWAS data is not yet available. Finally, we cannot interpret effect sizes in the same way as a clinical trial, as MR estimates reflect lifelong effects of “assignment” to genetic variants, not discrete interventions.

Overall, this study illustrates MR as a way to strengthen causal inference for the role of protective factors in mental health. Our results complement evidence from randomized controlled trials and prospective cohort studies indicating that physical activity has causal protective effects on depression (3–5). Drawing on instruments derived from large-scale cross-sectional GWAS of phenotypes can reduce the burden of conducting intensive randomized trials and/or collecting extensive exposure and outcome data over time. Further evidence of causal effects is of great importance because there are few known modifiable factors for depression. In conclusion, our findings validate physical activity as a causal protective factor for depression and point to the importance of objective measurement of physical activity in epidemiological studies in relation to mental health. Overall, this study supports the hypothesis that enhancing physical activity is an effective prevention strategy for depression.

## Acknowledgements and Funding

Dr. Choi was supported in part by a NIMH T32 Training Fellowship (T32MH017119). Dr. Smoller is a Tepper Family MGH Research Scholar and supported in part by the Demarest Lloyd, Jr. Foundation and NIH grant K24MH094614.

